# Notch signaling coordinates ommatidial rotation in the Drosophila eye via transcriptional regulation of the EGF-Receptor ligand Argos

**DOI:** 10.1101/728899

**Authors:** Yildiz Koca, Benjamin E. Housden, William J. Gault, Sarah J. Bray, Marek Mlodzik

**Affiliations:** Dept. of Cell, Developmental, and Regenerative Biology, Icahn School of Medicine at Mount Sinai, One Gustave L. Levy Place, New York, NY 10029; Graduate School of Biomedical Sciences, Icahn School of Medicine at Mount Sinai, One Gustave L. Levy Place, New York, NY 10029; Dept. of Physiology, Development and Neuroscience, University of Cambridge, Downing Street, Cambridge, CB2 3DY, UK

**Keywords:** Notch Signaling, Morphogenesis, Ommatidial Rotation, Photoreceptor Specification, *Drosophila*

## Abstract

In all metazoans, a small number of evolutionarily conserved signaling pathways are reiteratively used during development to orchestrate critical patterning and morphogenetic processes. Among these, Notch (N) signaling is essential for most aspects of tissue patterning where it mediates the communication between adjacent cells to control cell fate specification. In *Drosophila*, Notch signaling is required for several features of eye development, including the R3/R4 cell fate choice and R7 specification. Here we show that hypomorphic alleles of *Notch* – belonging to the *N*^*facet*^ class – reveal a novel phenotype: while photoreceptor specification in the mutant ommatidia is largely normal, defects are observed in ommatidial rotation (OR), a planar cell polarity (PCP)-mediated morphogenetic cell motility process. We demonstrate that during OR Notch signaling is specifically required in the R4 photoreceptor to upregulate the transcription of *argos (aos)*, an inhibitory ligand to the EGFR, to fine-tune the activity of Egfr signaling. Consistently, the loss-of-function defects of *N*^*facet*^ alleles and EGFR-signaling pathway mutants are largely indistinguishable. A Notch-regulated *aos* enhancer confers R4 specific expression arguing that *aos* is directly regulated by Notch signaling in this context via Su(H)- Mam dependent transcription.

## Introduction

*Drosophila* eye development serves as a paradigm for many developmental patterning processes and the dissection of the associated signaling pathways (Cagan and Ready, 1989a; Roignant and Treisman, 2009; Tomlinson and Ready, 1987; Wolff and Ready, 1991). The *Drosophila* eye consists of ~800 highly regularly arranged ommatidia, or facets, with each consisting of 8 photoreceptor (R-cell) neurons (R1-R8), arranged into a precise invariant trapezoidal pattern, and 12 accessory (cone, pigment, and bristle) cells (Tomlinson and Ready, 1987; Wolff and Ready, 1991). During larval stages, the eye develops from an imaginal disc, which is initially composed of identical pluripotent precursor cells. As a wave of cell proliferation and differentiation (referred to as morphogenetic furrow, MF) moves across the disc from posterior to anterior, it leaves regularly spaced preclusters of differentiating cells in its wake that will mature into ommatidia (Cagan and Ready, 1989a; Roignant and Treisman, 2009; Tomlinson and Ready, 1987; Wolff and Ready, 1991). At the 5-cell precluster stage, several patterning steps are apparent in addition to R-cell induction and differentiation, one being the differential specification of the two cells within the R3/R4 pair, which breaks the initial symmetry of the precluster. This differential R3/R4 specification requires the Wnt-Frizzled (Fz)/Planar Cell Polarity (PCP) pathway and its interplay with and asymmetric upregulation of Notch (N)-signaling (Blair, 1999; Cooper and Bray, 1999; Fanto and Mlodzik, 1999; Mlodzik, 1999; Strutt and Strutt, 1999). This cell fate induction step is followed by the rotation of the ommatidial precluster, referred to as ommatidial rotation, towards the dorsal-ventral (D/V) midline, the so-called equator (Mlodzik, 1999; Strutt and Strutt, 1999). As additional cells are recruited, the precluster undergoes a 90° rotation (in opposing directions in the dorsal and ventral halves of the eye) to establish the mirror-symmetric pattern most apparent in adult ommatidia along the D/V midline (Jenny, 2010) (see also Figure 1A-D).

**Figure 1.**
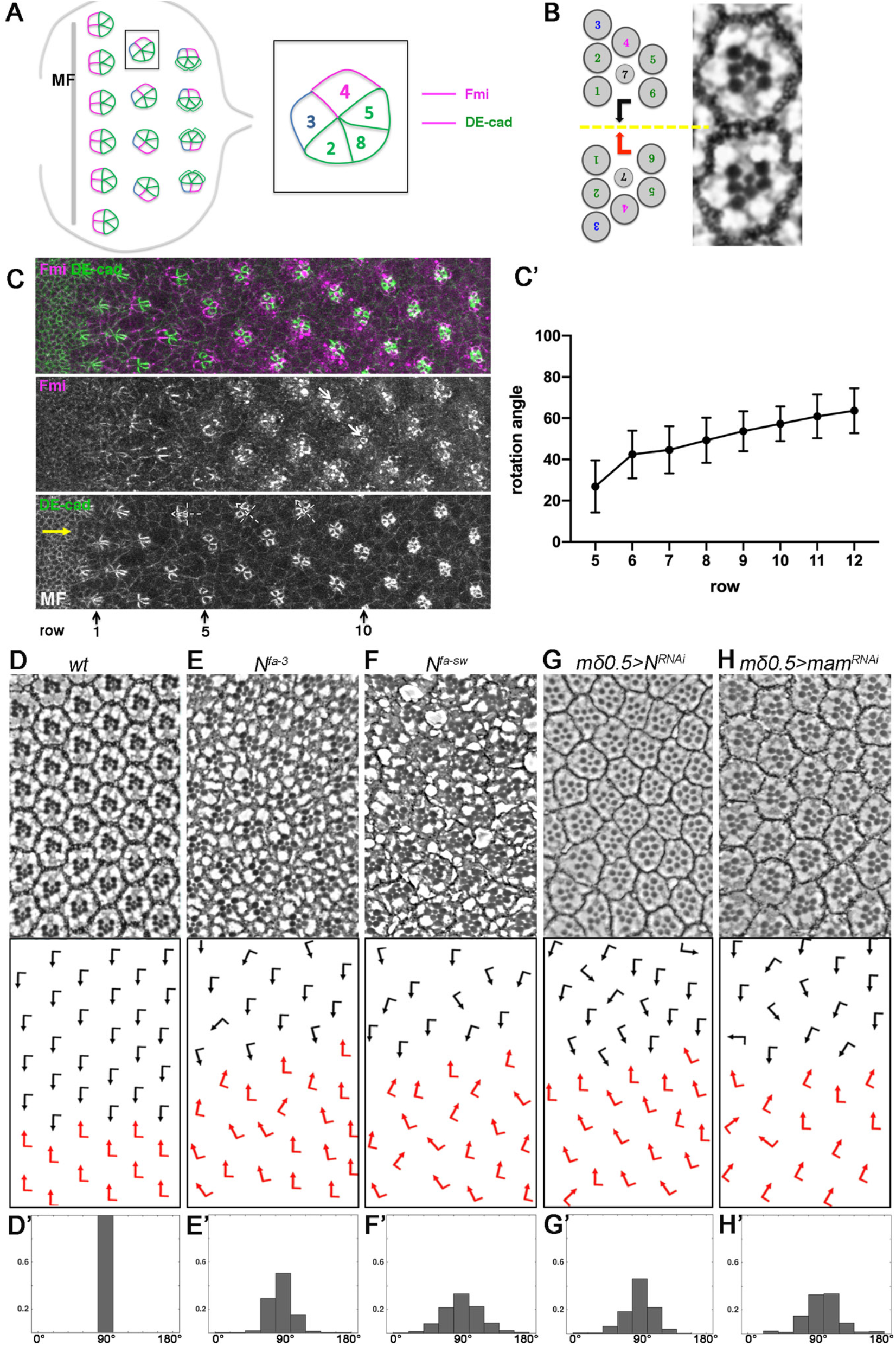
Perturbation of Notch signaling in the eye leads to misorientation of ommatidia. **(A)** Schematic of 3rd instar eye imaginal disc. As furrow (MF) moves across the eye disc from posterior to anterior ommatidial preclusters are forming in its wake, a process that involves lateral inhibition and R8 induction. R8 subsequently induces the sequential recruitment of R2/R5 and R3/R4 precursors pairs, resulting in the 5-cell precluster. Once the symmetry of 5-cell preclusters breaks due to differential R3/R4 specification, they start to rotate towards the the dorso-ventral midline (yellow line, “equator”) until they complete a 90° rotation and are aligned perpendicular to the equator. Fmi (magenta), initially detected in junctions of both R3/R4 precursors, becomes enriched to R4 junctional surfaces as the precursors mature. DE-cadherin (green) is upregulated in R2/R5 and R8 cells. **(B)** Schematic and section view of the two distinct chiral forms of adult ommatidia, displaying mirror image symmetry across the equator (yellow line). **(C)** Wild-type third larval instar eye imaginal disc stained for Fmi (magenta) and DE-cad (green) with MF at the anterior (left). Note junctional enrichment of Fmi in R4 (white arrows in Fmi monochrome). White dashed cross-arrows indicate orientation angle of preclusters. Yellow arrow marks position of the equator near MF. (**C’)** Quantification of OR angles at each row plotted for *wt* eye discs (45<n<60 per row, 8 eye discs). **(D-H)** Adult eye sections with orientation schematics (arrows are as in **B**). Note that the equator position is not affected. (**D’- H’**) Histograms of ommatidial orientation angles of respective genotypes shown in **D-H.** Wild type (*wt*) **(D, D’)**, *N^fa-3^* **(E, E’),** *N^fa-sw^* **(F, F’),** *mδ0.5*>*N*^*RNAi*^ *(BL7078*) **(G, G’)**, and *mδ0.5*>*mam*^*RNAi*^ (*BL63601)* **(H, H’);** n>300, 3 eyes per genotype.

Ommatidial rotation (OR) is a paradigm of PCP-mediated cell motility. Posterior to the MF, Wnt-Frizzled (Fz)/PCP signaling not only instructs the R3/R4 cell fate specification (Mlodzik, 1999; Strutt and Strutt, 1999), but also coordinates the direction and degree of OR. This is evident in core PCP mutants: e.g. *fz, flamingo/starry night (fmi/stan), or strabismus/Van Gogh (stbm/Vang*), which show defects in both R3/R4 specification (and hence ommatidial chirality) and the orientation of ommatidia (Das et al., 2002; Wolff and Rubin, 1998; Zheng et al., 1995). To date, several OR-specific regulators have been discovered based on the ommatidial misorientation phenotypes associated with their mutants (Brown and Freeman, 2003; Choi and Benzer, 1994; Chou and Chien, 2002; Fiehler and Wolff, 2007, 2008; Gaengel and Mlodzik, 2003; Mirkovic et al., 2011; Mirkovic and Mlodzik, 2006; Winter et al., 2001). For example, it is established that Fz/PCP signaling feeds into cadherin-based cell adhesion machinery through downstream effectors to precisely regulate the OR process (Mirkovic et al., 2011). EGF-Receptor (EGFR) signaling has also been shown to contribute to the process and genetic studies implicate input from EGFR signaling into cell adhesion factors (Brown and Freeman, 2003; Gaengel and Mlodzik, 2003). Genetic studies further suggest that cytoskeletal reorganization of ommatidial cells are coordinated with adhesion remodeling to drive the OR process downstream of Fz/PCP, EGFR, and potential other signaling pathways (Fiehler and Wolff, 2007; Gaengel and Mlodzik, 2003; Winter et al., 2001).

Notch (N) signaling is critical for cell fate determination in many if not all tissues in all metazoa mediating many essential cellular processes (Andersson et al., 2011; Bray, 2016). In particular in the *Drosophila* eye, Notch signaling is required at each step of eye development, ranging from the definition and growth of the eye field, to lateral inhibition within the MF to define correct precluster spacing, and to many aspects of cell fate induction of the individual R-cells and accessory cells including cone cell and pigment cell fate decisions (Doroquez and Rebay, 2006).

The widespread requirement in eye development means that many aspects of eye and ommatidial development are affected when Notch activity is perturbed (Cagan and Ready, 1989b; Papayannopoulos et al., 1998) causing a largely uninterpretable chaos and thus individual steps are very difficult to dissect.

Notch signaling initiates at the cell surface with the binding of a ligand (e.g. Delta) to the Notch receptor (Kopan and Ilagan, 2009; Rebay et al., 1991). Upon this interaction, the Notch receptor undergoes two sequential cleavages, releasing the Notch intracellular domain (NICD) and allowing for its translocation to the nucleus (Brou et al., 2000; De Strooper et al., 1999; Lieber et al., 2002; Mumm et al., 2000; Struhl and Greenwald, 1999). Once in the nucleus, NICD complexes with Mastermind (Mam) – a Notch pathway specific co-activator - and Suppressor of Hairless (Su[H]) - a transcription factor of the CSL family - to promote target gene expression (Bray, 2016; Struhl and Adachi, 1998; Wilson and Kovall, 2006; Wu et al., 2000). The precise signaling outcome depends on the genes that are regulated in each particular context (Bray, 2016; Bray and Gomez-Lamarca, 2018). Given the multiple requirements for Notch activity in eye and ommatidial development (Cagan and Ready, 1989b; Papayannopoulos et al., 1998), it is likely that different primary targets will be involved in implementing each distinct and individual role.

A well-established function of Notch signaling is in the context of R3/R4 cell fate specification in cooperation with Fz/PCP signaling (Cooper and Bray, 1999; Fanto and Mlodzik, 1999; Tomlinson and Struhl, 1999). At the 5-cell precluster stage, among the R3/R4 precursors, the cell that is closer to the equator ends up having higher Fz/PCP signaling activity, specifying it as an R3 cell. Fz-PCP signaling-dependent transcriptional upregulation of *Delta* (*Dl*) and *neuralized* (*neu*) in R3 subsequently induces the adjacent cell of the pair as R4 by a classical Dl-N interaction, thus activating the Notch pathway to higher levels in the R4 precursor (Cooper and Bray, 1999; del Alamo and Mlodzik, 2006; Fanto and Mlodzik, 1999; Tomlinson and Struhl, 1999; Weber et al., 2000). Since N-signaling is critical for R3/R4 asymmetry, its perturbation might also cause OR phenotypes brought about by R4 specification defects. However, strikingly, a class of hypomorphic *N* alleles, the so-called *facet* alleles *N*^*fa-3*^ and *N*^*fa-sw*^ (Markopoulou et al., 1989), exhibits misorientation of ommatidia yet largely normal eye patterning with correct R4 specification and ommatidial chirality establishment (Figure 1E,F). This orientation-specific ommatidial phenotype is reminiscent of previously identified mutants that are linked to OR (e.g. *nmo* or *argos*^*rlt*^ (Choi and Benzer, 1994; Gaengel and Mlodzik, 2003) and suggests that *Notch* signaling has a direct role in rotation which is independent of R4 specification *per se*. A potential role of Notch signaling in OR and associated morphogenesis, however, has not yet been explored.

Using a combination of phenotypic analyses, genetic interactions, cell-based and molecular studies, we define here an OR-specific role for Notch signaling after R4 fate specification, in addition to its well described function in cell fate choices in ommatidial patterning. We demonstrate that Notch signaling coordinates the morphogenetic process of OR by fine-tuning the activity of the EGFR pathway and PCP signaling. Specifically, Notch signaling in R4 leads to a direct transcriptional upregulation of *argos* (*aos*), as confirmed by Su(H) DNA occupancy and reporter expression studies. As Argos is an inhibitory ligand to EGFR and EGFR-signaling is required for OR regulation, the loss or reduction of Notch-dependent *aos* expression leads to an imbalance between positively and negatively acting EGFR ligands and hence to defects in the OR outcome. In addition, Notch signaling affects the levels of the PCP protein Flamingo (Fmi) in R4. This dual Notch signaling input into EGFR and PCP-pathways orchestrates a precise OR process and hence demonstrates a critical and specific role of Notch activation during OR.

## Results

### Notch signaling is required in R3/R4 pairs for accurate ommatidial rotation

Ommatidial rotation (OR) is a morphogenetic process that occurs during larval eye development, and it results in the final orientation of ommatidia, forming a mirror image arrangement across the dorso-ventral (D/V)-midline in the adult eyes (Figure 1A-D). OR is instructed by Fz/PCP signaling and associated pathways including Notch and EGFR-signaling that are involved in R3/R4 photoreceptor fate specification (Brown and Freeman, 2003; Cooper and Bray, 1999; Fanto and Mlodzik, 1999; Gaengel and Mlodzik, 2003; Strutt and Strutt, 2003; Tomlinson and Struhl, 1999).

At larval stages, as the morphogenetic furrow (MF) sweeps across the eye disc form posterior to anterior, it induces the formation of a new row of regularly spaced ommatidial preclusters every ~2 hours (Fig. 1A,C, and (Tomlinson and Ready, 1987; Wolff and Ready, 1991), giving rise to rows of ommatidial clusters that are ~2 hours apart from each other in developmental time and thus allowing the visualization of progressively more mature clusters in the same tissue sample (Fig. 1A,C; also (Jenny, 2010). As such, these rows reflect consecutive stages of ommatidial maturation allowing for the tracking of OR row by row as the eye disc develops (Fig. 1C-C’). The use of apical junctional markers, like E-cadherin (E-cad, enriched at the junctions of R2/R5 and R8 cell boundaries), and Flamingo (Fmi, enriched at the apical junctions of R4) allows for the tracking of OR angles of individual clusters during development (Fig. 1A,1C-C’). In wild type, ommatidial (pre)clusters initiate rotation in row 5 and largely complete the process by rows 14- 15 (Figure 1C-C’), resulting in a 90° rotation angle that aligns mature ommatidia perpendicular to the D/V midline in the adult (Figure 1B,1D-D’; note that the final angle in wild-type is an invariant 90°, Fig. 1D-D’; (Jenny, 2010).

To investigate the role of Notch signaling in OR, we first analyzed two recessive hypomorphic *Notch* (*N*) alleles: *facet-strawberry* (*N*^*fa-sw*^) and *facet-3* (*N*^*fa-3*^). The *facet* class of a *Notch* alleles have been characterized and are thought to be caused by either an insertion\s of transposable element into an intronic region of *Notch* or deletion of non-transcribed sequences in the locus, thus not affecting the coding sequence and ultimate protein product but causing a reduction in gene expression in certain contexts (Markopoulou et al., 1989). In comparison to wild type, hemizygous *N*^*fa-sw*^ and *N*^*fa-3*^ males showed frequent misrotations of ommatidia, including both under- and over-rotation of individual clusters. Surprisingly, besides the OR defects, these mutant eyes were normal in eye size, photoreceptor specification, chiral arrangements, and other aspects of eye development (Figure 1E-F’; also (Markopoulou et al., 1989).

Since differential R3/R4 cell specification is critical for OR (Mlodzik, 1999; Strutt and Strutt, 1999), we next asked if Notch signaling is required in this cell pair for the OR process. To separate the potential role of Notch signaling in OR from its role in the R3/R4 cell fate choice, we took advantage of the Gal4/UAS-system (Brand and Perrimon, 1993), which allows for temporal knock-down of the respective genes and employed a driver (*mδ0.5-Gal4*) that is active in the R3/R4 pair and is upregulated in R4, as a result of *Notch*-mediated R4 specification (Cooper and Bray, 1999), where it is subsequently maintained during the OR process. As it is up-regulated in response to R4 specification, gene targeting with this driver should not significantly affect cell fate specification within the R3/R4 pair. Knockdown of *Notch* via *mδ0.5-Gal4* in R3/R4 cells phenocopied the misorientation phenotype(s) observed in the *N*-*facet* alleles, without affecting R3/R4 cell fate choices, suggesting that *Notch* activity is required in R3/R4 cells, and predominantly in R4, after cell fate determination to regulate OR (Figure 1G-G’, Figure S1A-A’). To ask whether this requires Notch-mediated transcriptional activation, we tested the requirement for *mastermind* (*mam)*, a Notch-specific transcriptional co-activator (Wu et al., 2000). As the Notch-associated DNA binding factor, *Su(H),* is also required for transcriptional repression, its perturbation could yield complex effects making it a less suitable choice (Barolo et al., 2002; Furriols and Bray, 2000; Morel et al., 2001; Wu et al., 2000). Strikingly, when knocked-down in the same *mδ0.5-Gal4* based assay (*mδ0.5*>*mam*^*RNAi*^), depletion of *mam* produced very similar phenotypes to the *N* knock-down, displaying OR defects with over- and under-rotated ommatidia (Figure 1H-H’, Figure S1B-B’) thus suggesting that Notch-dependent transcription is critical for accurate OR.

### Notch signaling affects OR during the developmental process in eye imaginal discs

To rule out the possibility that the misorientation phenotypes observed in adults upon *Notch* signaling perturbation were due to a later secondary cell packing effect – Notch is required for several late steps in ommatidial patterning (Cagan and Ready, 1989b)- we followed the rotation of ommatidial clusters during development in eye discs at the time of the OR process (see for example Figure 1C for wild-type). As a standard clonal analysis cannot be used for Notch pathway components due to the multitude of steps affected by Notch signaling in the eye disc (Cagan and Ready, 1989b), we again used the *mδ0.5-Gal4*-mediated knockdown (KD) strategy. In contrast to wild-type eye discs, KD of *Notch* or *mam* in R4 cells led to an abnormal rotational pattern, where ommatidial preclusters displayed a significantly wider range of rotation angles as compared to wild-type (Figure 2, and Figure S2), consistent with the under and over-rotation phenotypes seen in adult eyes (Fig. 1G-H’). Taken together, these data indicate that Notch signaling controls, via its transcriptional activation function, the rotation of ommatidial clusters during early stages of the process during eye development.

**Figure 2.**
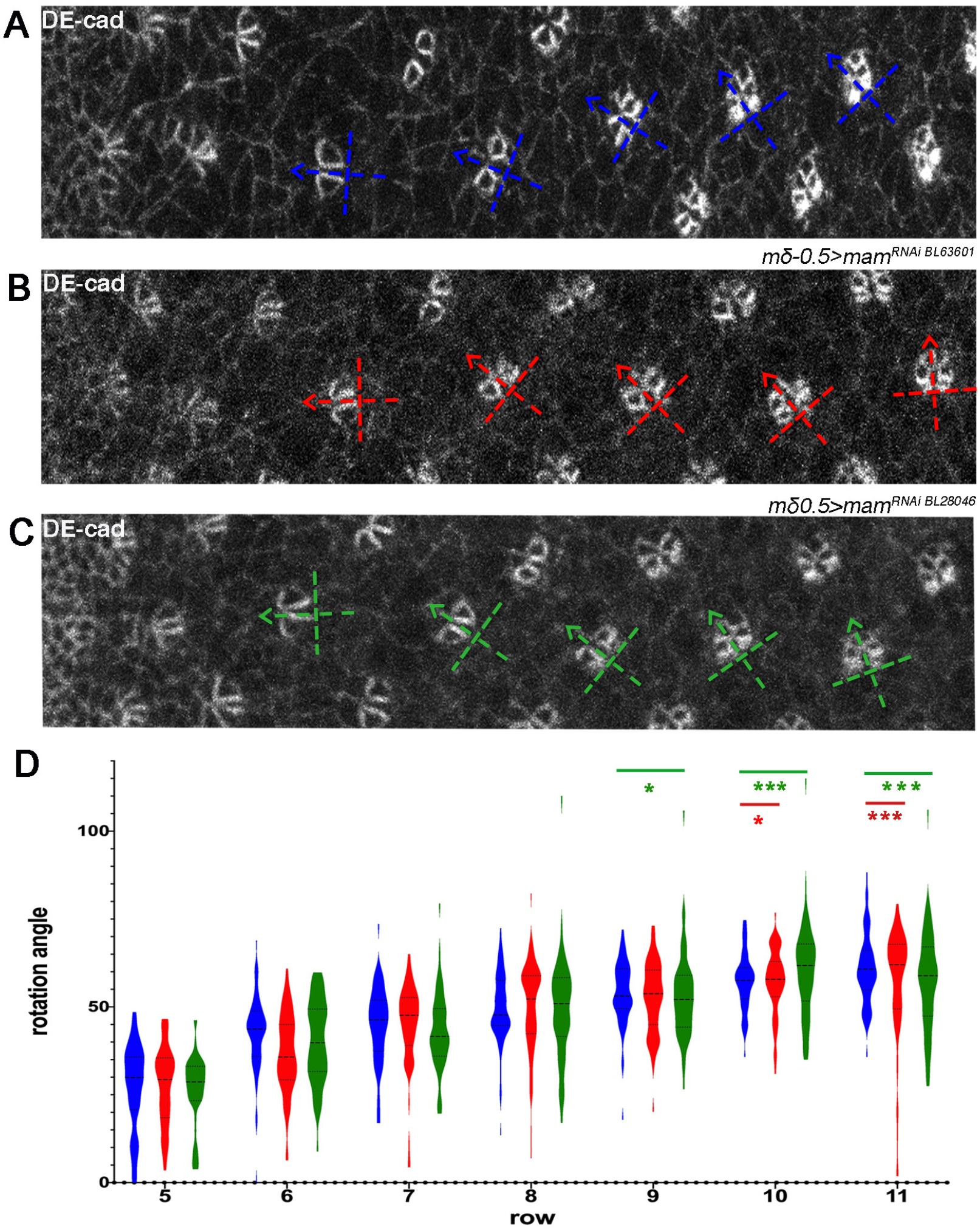
Notch signaling is required in R3/R4 pairs to regulate OR. **(A-C)** Third instar larval eye imaginal discs stained for DE-cad in *wild type* **(A)**, *mδ0.5*>*N*^*RNAi*^ *(BL31383)* **(B)**, and *mδ0.5>mam^RNAi^ (BL28046*) **(C)**. Blue, red and green dashed cross-arrows, respectively, indicate the orientation of ommatidial preclusters for each genotype. **(D)** Quantification of rotation angles observed in individual preclusters in rows 5-11, plotted for control (*wild type*) in blue; *mδ0.5*>*N*^*RNAi*^ (red); and *mδ0.5*>*mam*^*RNAi*^ (green). Statistical analyses were performed for each row between control (blue) and colored genotypes. Asterisks denote significance by chi-square test (* *p*<0.05, ** *p*<0.005, *** *p*<0.0005). Note the generally wider spread of rotation angles in most rows in the experimental genotypes as compared to wild-type. Scale bars indicate 10 μm.

### Notch signaling genetically interacts with *argos* in OR establishment

To get further insight into how Notch signaling might mechanistically affect OR, we tested for genetic interactions between the *mδ0.5*>*mam*^*RNAi*^ genotype and known regulators of OR. Several genes have been implicated in regulating OR based on ommatidial misrotation observed in their mutants and the role of several of these has been further validated by functional and molecular studies (Brown and Freeman, 2003; Choi and Benzer, 1994; Chou and Chien, 2002; Cooper and Bray, 1999; Fiehler and Wolff, 2007, 2008; Winter et al., 2001).

We used a *mδ0.5*>*mam*^*RNAi*^ combination that caused mild OR defects at low temperatures and asked whether its phenotype can be dominantly modified by other OR-associated genes. Among the known OR regulators, we detected a specific interaction with multiple alleles of *argos* (*aos*; including a deficiency for the gene), whereas other OR-associated genes tested did not show an interaction (Figure 3, Suppl. Data, Table S1). In parallel, we also asked whether the core PCP genes could modify the *mδ0.5*>*mam*^*RNAi*^ OR phenotype, as PCP factors contribute to OR (Das et al., 2002; Wolff and Rubin, 1998; Zheng et al., 1995). Whereas most core PCP genes did not dominantly affect the *mδ0.5*>*mam*^*RNAi*^ phenotype, alleles of *prickle* (*pk)* did enhance the OR defects (Suppl. data, Figure S3). Neither *aos* nor *pk* heterozygosity affected OR or other aspects of eye development on its own, confirming that their interaction with the *mδ0.5*>*mam*^*RNAi*^ background is not an additive feature (Figure S4). The *pk*^−/+^ effect was surprising, because *pk* itself does not display much of an OR phenotype in comparison to the other core PCP genes. However, *pk* is genetically required in R4 (Jenny et al., 2003), where *mδ0.5-Gal4* is driving expression and so an R4 specific interaction could be envisioned (see Discussion).

**Figure 3.**
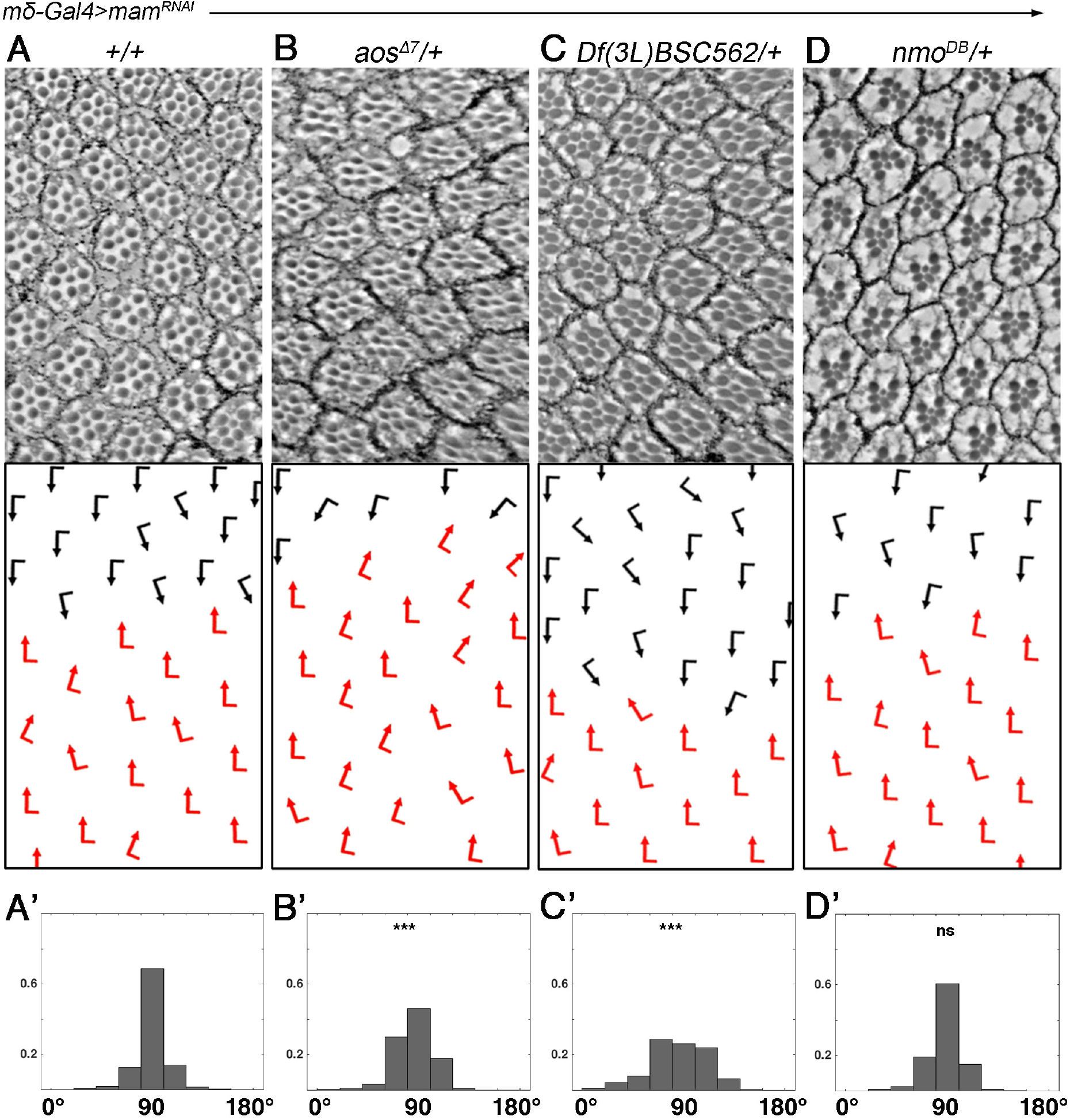
*mam* genetically interacts with *aos* during the OR process. **(A-D’)** Adult eye sections with ommatidial orientation schematics (arrows as in Figure 1) and orientation angle histograms of eyes of the genotypes indicated. All genotypes are *mδ0.5Gal4*, *UAS-mam*^*RNAi*^ *(BL28046) mδGal4*>*mam*^*RNAi*^ in the following genetic backgrounds: **(A-A’)** +/+ (*wild-type* control); **(B-B’)** *aos^Δ7^*/+, **(C-C’)** *Df(3L)BSC562*/+ (deletion of *aos* gene), and **(D-D’)** *nmo*^*DB*^/+. Asterisks denote significance by chi-square test (\*\*\**p*<0.0005). Note robust enhancement of the *mδ0.5Gal4*>*mam*^*RNAi*^ rotation phenotype by both *aos*^−/+^ genotypes. See supplemental material for additional genotypes.

Taken together, these genetic data raised the possibility that Notch signaling modulates the OR process via the EGFR pathway, because Argos is a secreted EGFR ligand that inhibits the receptor function (Klein et al., 2004; Klein et al., 2008; Vinos and Freeman, 2000), and via R4-associated PCP signaling. As the enhancement(s) are associated with a reduction in the transcriptional output of Notch signaling, caused by the *mδ0.5*>*mam*^*RNAi*^ genotype, the expression of some of these genes might be directly regulated by Notch-signaling in R4.

### Notch promotes *aos* expression in R4

To determine how Notch signaling interacts with *aos* during OR, we next examined the expression pattern of *aos* during eye disc patterning and development. Several *aos* alleles have been associated with ommatidial misrotation, notably *roulette* (*aos*^*rlt*^) was identified as an OR specific mutation (Choi and Benzer, 1994), even before *aos* was characterized as an EGFR ligand. Aos was subsequently used as a tool to define the role of EGFR/Ras signaling in OR (Brown and Freeman, 2003; Gaengel and Mlodzik, 2003; Strutt and Strutt, 2003). Interestingly, although *aos* is expressed at base levels in all photoreceptors of developing ommatidia (Figure 4A, also (McNeill et al., 2008; Yan et al., 2009), it was specifically upregulated in the R4 cell during the OR process, as detected by an enhancer trap reporter for the gene (*aos-lacZ*; Figure 4A-A’). We thus asked whether this *aos* upregulation is dependent on Notch/Mam-signaling. To this end, we employed *mδ0.5*>*mam*^*RNAi*^ in a mosaic clonal manner, which allows for direct comparison of wild-type and *mam* KD ommatidia within the same tissue. In this assay, *mam* KD caused a marked reduction in *aos-lacZ* expression levels in R4 (Figure 4B,D). This is consistent with the hypothesis that Notch signaling activation in R4 is required for *aos* upregulation in this cell. The levels of Elav, a nuclear neuronal (all R-cells) marker, were unchanged in the same genetic scenario (Figure 4F, also Suppl. data Figure S5), indicating that Notch signaling activity in R4 specifically affects *aos* transcription.

**Figure 4.**
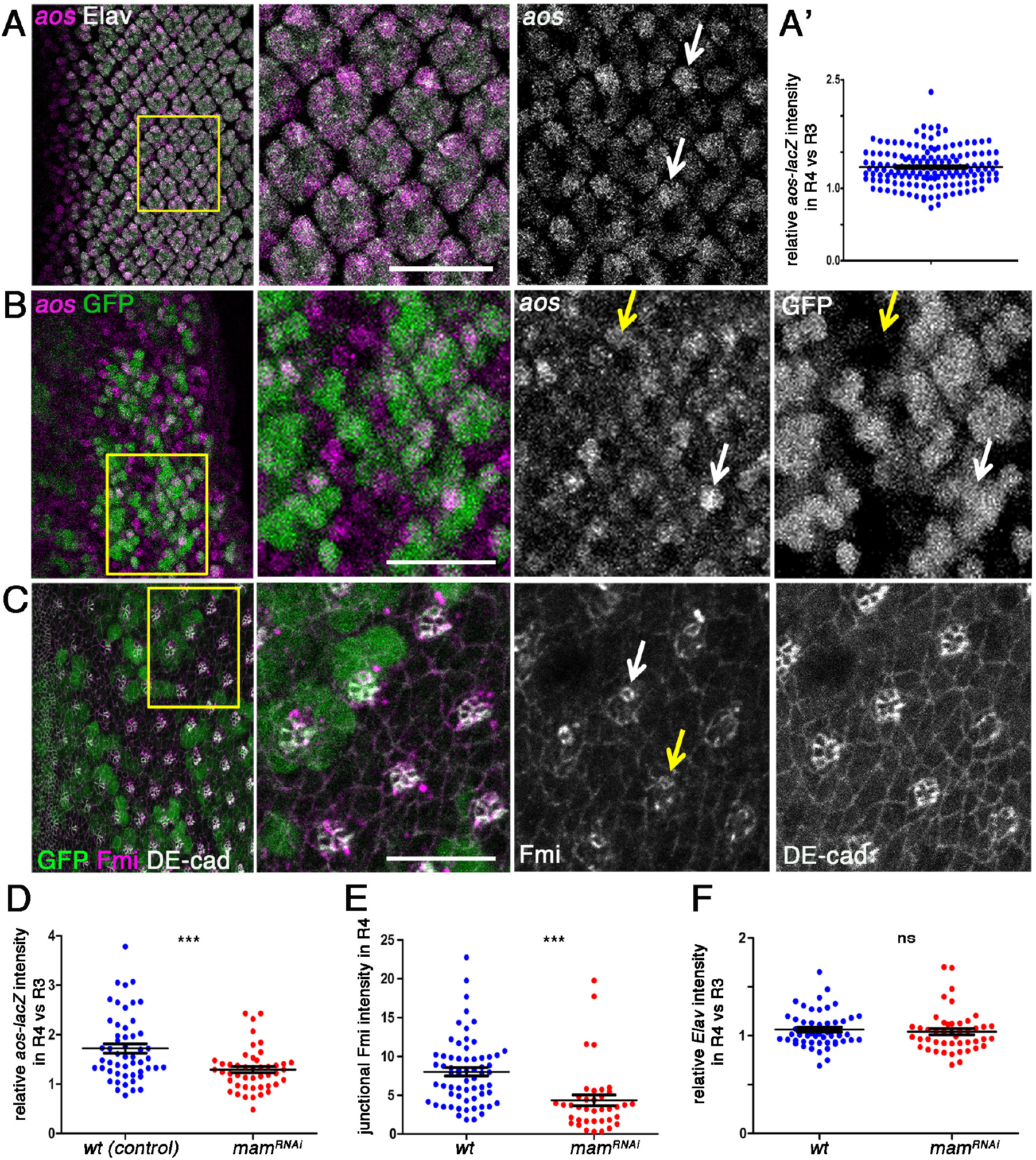
Notch signaling promotes *aos* expression (*aos-lacZ*) and junctional Fmi enrichment in R4. **(A)** Wild type third larval instar eye imaginal disc stained for Elav (gray) and *aos*-lacZ (magenta and monochorme panel). (**A’**) Quantifcation of expression level of aos-lacZ in R4 relative to R3 (see also Supplemental data for control quantification fo Elav). (**B-C**) Third larval instar eye imaginal discs mosaic for *mδ0.5*>*mam*^*RNAi*^ (*BL28046*; marked by absence of GFP/green) stained for *aos-lacZ* (magenta in **B** and monochrome); Fmi (gray in **C** and monochrome) and DE-cad (magenta in **C** and monochrome). White and yellow arrows point at R4 cells in wild type and mutant tissue, respectively; note reduction of *aos-lacZ* and Fmi staining in mutant areas of the respective panels. (**D**) Quantification *aos-lacZ* expression in R4 relative to R3 plotted for individual clusters in *wt*-control (blue) and *mδ0.5*>*mam*^*RNAi*^ (red). **(E)** Quantification of junctional Fmi intensity normalized to DE-cad staining in R4 plotted for individual clusters for *wt* (blue) and *mδ0.5*>*mam*^*RNAi*^ (red). **(F)** Quantification of Elav intensity in R4 relative to R3 plotted for individual clusters with wt (blue) and *mδ0.5*>*mam*^*RNAi*^ (red). Asterisks denote significance by chi-square test (*** *p*<0.0005). Scale bars indicate 10 μm.

Upon *Notch*-mediated R4 cell specification, several of the core PCP factors are enriched in R4. This is most evident with the increase of Flamingo (Fmi, also called *starry night/stan*) levels at the apical junctional region in R4. Fmi is an atypical cadherin that plays a central role in PCP establishment by stabilizing the core PCP complexes at junctional regions across cell membranes (Wu and Mlodzik, 2009). In particular in eye discs, before R3/R4 differentiates, Fmi is apically enriched in both precursors. As the symmetry of the precluster breaks and R4 is specified, it becomes enriched in the apical surface of R4 (Das et al., 2002) and this upregulation serves as an R4 marker (see also Figure 1A,C). In *fmi* mutants, ommatidia adopt a random chiral form or lose asymmetry, and additionally display misrotation defects (Das et al., 2002). As we detected a genetic interaction between *mδ0.5*>*mam*^*IR*^ and *pk*^−/+^, we also examined Fmi expression as an indicator of core PCP factor levels. Upon comparing the Fmi expression pattern/junctional levels between *mam* KD and neighboring wild-type ommatidia in the respective mosaic eye discs (employing again mosaic clonal *mδ0.5*>*mam*^*RNAi*^-mediated KD), we observed a significant reduction in apical Fmi levels in Mam-depleted R4 cells (Figure 4B,D) suggesting a Notch-mediated upregulation of core PCP factors in R4 (see below and Discussion).

### *argos* is a direct R4-specific transcriptional target of Notch signaling

To corroborate and refine the hypothesis that Notch-signaling directly regulates the transcription of *aos*, we examined whether the *aos* locus was occupied by with the core Notch-associated transcription factor, Su(H), in a genome-wide chromatin immunoprecipitation (ChIP) data-set from *Drosophila* larval central nervous system (Zacharioudaki et al., 2016). Strikingly there was significant enrichment of Su(H) within the first intron of *aos* overlapping with predicted conserved Su(H) binding-motifs, consistent with a direct regulation by Notch/Mam/Su(H) complexes (Fig. 5A). To ask which cells in the developing eye disc are susceptible to this regulatory input, we utilized a GFP reporter construct, encompassing the high confidence “peak” region from the Su(H) ChIP, which had previously been shown to respond to Notch in muscle progenitor cells (*aos1-GFP*, Fig. 5B; (Housden et al., 2014) and tested its expression in the eye discs. Strikingly, expression of *aos1-GFP* was detected predominantly in R4 cells, as confirmed by co-expression of the R4 marker *mδ0.5-lacZ* (Figure 5C). The transcriptional regulation of *aos* by Notch/Mam-Su(H) in R4 appears specific, as there was little or no Su(H) binding detected at the *fmi* or *pk* loci in the same ChIP data-set (Suppl. Figure S6). Taken together, these data indicate that *aos* is a specific transcriptional target of Notch-signaling in R4, and that the visible increase in Fmi protein in R4 is likely due to other post-transcriptional mechanisms (see Discussion).

**Figure 5.**
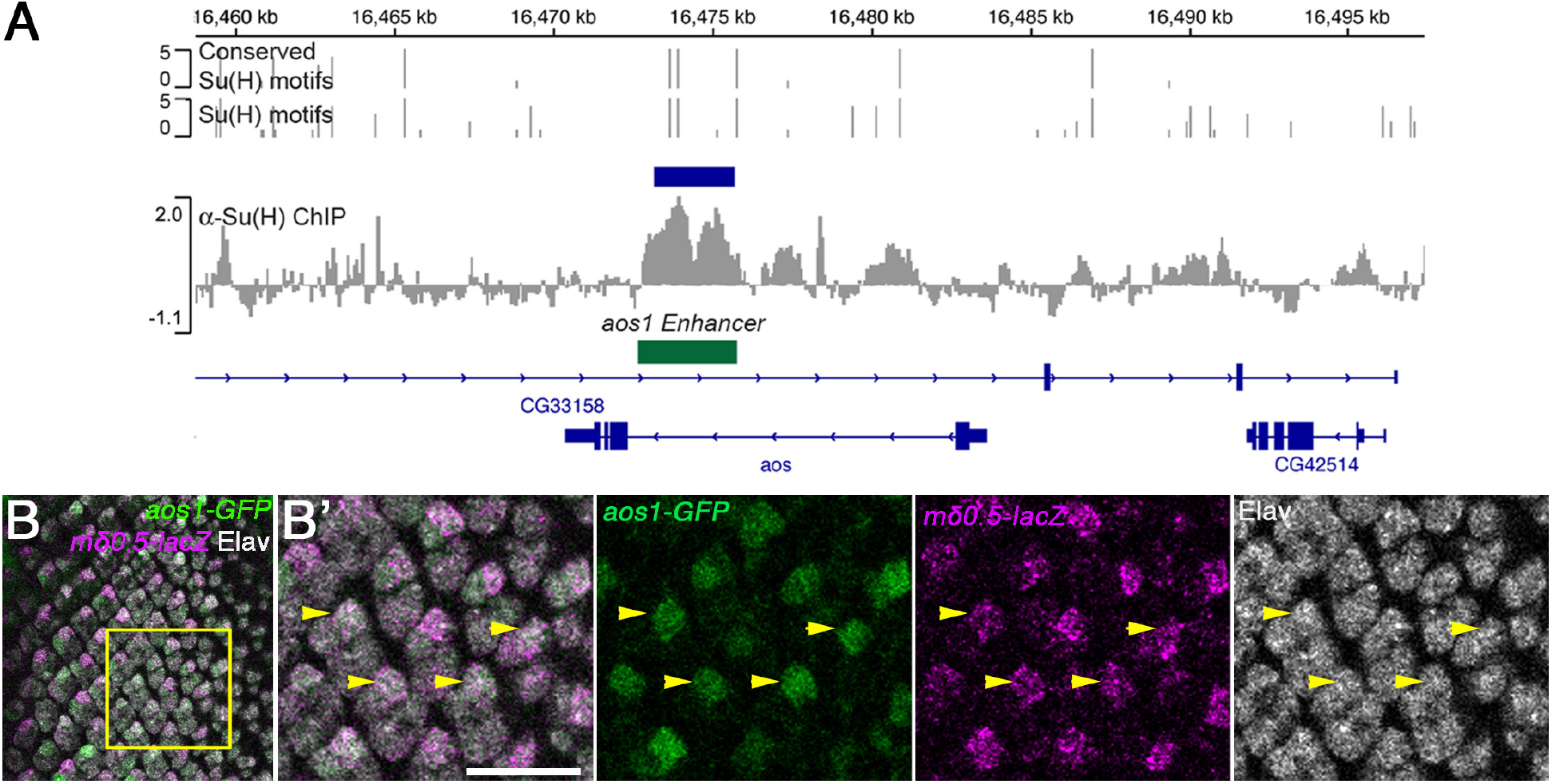
Notch-Su(H) signaling directly promotes *aos* expression in R4. **(A)** ChIP enrichment for Su(H)-occupancy at the *aos* locus in CNS samples (α-Su(H) enrichment relative to input, scale log_2_) (Zacharioudaki et al., 2016). Blue bar indicates the region of significant enrichment. Gray bars indicate the positions of Su(H)-binding motifs; bar height represents the motif-score (scale 0-5); upper graph indicates motif conservation across 12 *Drosophila* species. Gene regions are depicted in dark blue. Green bar indicates the genomic region used in the *aos1-GFP* reporter construct (Housden et al., 2014). **(B-B’)** Third instar eye imaginal disc stained for *aos1-GFP* (green), Elav (nuclei of all R-cells, gray), and *mδ0.5-lacZ* (magenta, R4-specific marker). **B’** panels show higher magnification of boxed area in B. Yellow arrowheads highlight examples of R4 cells revealing the co-expression of *aos1-GFP* and the R4 marker *mδ0.5-lacZ*. Scale bar indicates 10 μm.

## Discussion

The involvement of Notch signaling in controlling cell proliferation, cell differentiation, and patterning has been studied in a vast set of contexts ranging from neuronal development to intestinal homeostasis in flies and vertebrates (Andersson et al., 2011; Bray, 2016; Henrique and Schweisguth, 2019). Spatial and temporal control of Notch activity, along with the employment of cell/tissue specific downstream elements and crosstalk with other signaling pathways, confers the functional versatility and specific reiterative use of Notch pathway activation (Andersson et al., 2011; Bray, 2016; Doroquez and Rebay, 2006; Guruharsha et al., 2012; Henrique and Schweisguth, 2019). In the *Drosophila* eye alone, for instance, Notch signaling has a defined and specific function at nearly every stage of tissue development and patterning (Doroquez and Rebay, 2006). At early larval stages Notch activity in the eye is restricted to the dorsoventral equator, from which it promotes the growth of the eye disc and eye field within the disc, and the formation of the MF (Dominguez and de Celis, 1998; Kenyon et al., 2003; Reynolds-Kenneally and Mlodzik, 2005). Within and posterior to the MF, Notch signaling is first required for the spacing of the R8 precursors and thus ommatidial preclusters (Baker et al., 1990), and the subsequent specification of R3/R4 and R7 fates in a stepwise fashion (Baonza and Freeman, 2001; Cooper and Bray, 1999, 2000; Fanto and Mlodzik, 1999; Tomlinson and Struhl, 1999, 2001). As ommatidial clusters mature further, Notch signaling controls the acquisition of the cone and pigment cell fates and apoptosis of the non-committed remaining interommatidial cells to generate the precise and highly ordered pattern of a fully-developed ommatidia (Cagan and Ready, 1989b).

At each stage, Notch signaling acts in concert with multiple other pathways in a spatially and temporally restricted manner in order to achieve particular and specific readouts. For example, Fz/PCP signaling pathway triggers Delta expression in R3 to induce Notch activation and the resulting R4 cell fate in the adjacent cell of the R3/R4 pair (Cooper and Bray, 1999; Fanto and Mlodzik, 1999; Weber et al., 2000). Furthermore, at nearly every step mentioned, there is cross-talk between Notch and EGFR signaling pathways (including the R3/R4 specification steps; (Weber et al., 2008) to achieve the respective developmental outcome (Doroquez and Rebay, 2006). However, the nature of the interaction between Notch and EGFR pathways, and the downstream elements engaged, differ depending on the context. For example, in the course of R7 specification, Notch promotes the expression of the transcriptional repressor Yan which in turn needs to be post-translationally repressed by EGFR signaling to establish R7 fate (Rohrbaugh et al., 2002). On the other hand, Notch and EGFR signaling effectors combinatorially drive the expression of the *Drosophila* paired box gene 2 (dPax2) to promote cone cell identity (Flores et al., 2000). The interactions of Notch signaling with the EGFR and the Wnt-Fz/PCP pathways have not only been well-documented in eye patterning but also in other developmental contexts and cancer, highlighting the importance of the communication of Notch with the respective pathways during development and disease (Baker et al., 2014; Bao, 2014; Capilla et al., 2012; Haruki et al., 2005; Hasson and Paroush, 2006; Yoo et al., 2004).

Our results document a new function for Notch signaling in R4 to govern the morphogenetic process of OR, which is independent of its role in R4 cell fate acquisition. Perturbation of Notch signaling pathway components in the R3/R4 pair, and R4 in particular, after the cell fate choice is established, leads to the misregulation of the rotation process. In this context, Notch signaling regulates *aos and fmi, as* their expression levels in R4 are diminished upon downregulation of Notch signaling. Our data demonstrate that *aos* transcription is directly regulated by Notch, via Su(H)/Mam/NICD mediated transcriptional control, but the effect on *fmi* levels is less well defined, as there is no clear evidence to suggest direct transcriptional regulation in this case. The effect of Notch on apical Fmi levels is unlikely to be a secondary effect of *aos* deregulation, since Fmi has been reported to be expressed in R4 and ectopically in R3 in the absence of *aos* (Gaengel and Mlodzik, 2003). Yet, how Notch-signaling in R4 regulates the levels and function of the core PCP factors in R4 remains largely elusive at this time.

Based on previous reports in *Drosophila*, the effect of Notch signaling on *aos* expression is context dependent (Housden et al., 2014). Gene expression is often controlled by multiple *cis*-regulatory units that integrate information from various transcriptional inputs. Essentially, *aos* has been reported to exhibit context-dependent enhancer selection in the wing: *aos* contains three enhancer regions identified, that are differentially responsive to factors that act downstream of Notch or EGFR signaling pathways (Ajuria et al., 2011; Krejci et al., 2009). The presence of context-determining factors will determine whether an enhancer is primed to react to a specific signal, which may result in a gene having different responses towards the same signals depending on which enhancer is accessible (Housden et al., 2014). Notably, the *aos1-GFP* reporter experiment reported here, and the Su(H) ChIP data, argue that the *aos1* enhancer is directly responsive to Notch signaling in R4, indicating that Notch activates *aos* expression in this context.

Variations in the phenotypes from mutations in rotation-specific genes are indicative of their function in OR. For example, mutations disrupting the *nmo* kinase result in a severe under-rotation of ommatidia arguing that it has a positive role in promoting rotation (Fiehler and Wolff, 2008; Mirkovic et al., 2011). In contrast, *aos*^*rlt*^ mutants exhibit random rotation angles with both under- and over-rotated ommatidia (Brown and Freeman, 2003; Gaengel and Mlodzik, 2003; Strutt and Strutt, 2003). This suggests that the consequent change in EGFR signaling results in an overall misregulation of rotation, rather than promoting or inhibiting ommatidial motility *per se*. Consistent with the notion that Notch-signaling in R4 directly regulates *aos* expression, the phenotype caused by *facet* alleles of *Notch* largely mimics that of *aos*^*rlt*^, with an overall deregulation of the process and resulting random rotation angles. All R3/R4-specific Notch or Mam RNAi interference scenarios also mimic these phenotypes. Given that OR entails the coordination of cytoskeletal and adhesion dynamics (Chou and Chien, 2002; Fiehler and Wolff, 2007; Gaengel and Mlodzik, 2003; Mirkovic et al., 2011), our data suggest an input from Notch signaling into these molecular processes through negative regulation of EGFR-signaling and possibly also its interplay with PCP signaling.

In recent years, involvement of Notch signaling in morphogenesis has been suggested in various contexts, including *Drosophila* oogenesis, zebrafish sensory organ development, and human vascular barrier formation (Dobens and Raftery, 2000; Kozlovskaja-Gumbriene et al., 2017; Torres et al., 2003). These studies also suggest an input from Notch signaling into the cell adhesion and/or cytoskeletal factors, mostly through Notch-mediated transcription of genes that regulate adhesion and cytoskeletal dynamics (Pezeron et al., 2014) although a direct input from the Notch receptor into adhesion has also been revealed (Polacheck et al., 2017). Overall, the multifaceted involvement of Notch signaling in cellular (re)organization and morphogenesis is becoming increasingly evident. Future studies will be needed to provide insight into the mechanistic details of how Notch can mediate distinct morphogenetic processes. As Notch signaling has long been implicated in cancer metastasis, such studies will also hold promise for better understanding of disease and hence future therapeutic applications.

## Materials and Methods

### Fly strains and genetics

Flies were raised on standard medium and maintained at 25°C unless otherwise stated.

*N*^*fa-sw*^ and *N*^*fa-3*^ were gifts from Spyros Artavanis-Tsakonas.

*nmo*^*DB*^/*TM6b and mδ0.5-Gal4 FRT40/SM3:TM6b* were from Mlodzik lab stocks.

*aos*^*Δ7*^/*TM3*, *Df(3L)BSC562/TM3*, *pk*^*pk-sple13*^/*CyO*, *pk*^*pk-sple6*^/*CyO*, *w*^*1118*^, *Notch*^RNAi^ (BL31383, BL7078) and *mam*^*RNAi*^ lines (BL28046, BL63601) were ordered from Bloomington Drosophila Stock Center.

*aos-lacZ/TM6b* was a kind gift from Utpal Banerjee.

*aos1-GFP/TM3* was from Bray lab stocks (Housden et al., 2014).

*mδ0.5*>*N*^*RNAi BL31383*^ (*mδ0.5-Gal4/*+; *UAS-N*^*RNAi BL31383*^/+) were obtained at 25°C.

*mδ0.5*>*mam*^*RNAi BL28046*^ (*mδ0.5-Gal4*; *UAS-mam*^*RNAi BL28046*^ /+) were obtained at 18°C.

*mδ0.5*>*N*^*RNAi BL7078*^ (*mδ0.5-Gal4*/+; *UAS-N*^*RNAi BL7078*^/+) were obtained at 18°C.

*mδ0.5*>*mam*^*RNAi BL63601*^ (*mδ0.5-Gal4*, *UAS- mam*^*RNAi BL63601*^/+) were obtained at 18°C.

Control eye disc stainings were done in *mδ0.5-Gal4 FRT40*/+ background.

Genetic interactions were tested at 25°C between *mδ-Gal4/+; UAS-mam^RNAi BL28046^*/+ and the heterozygosity of the respective genes.

*mδ0.5*>*mam*^*RNAi BL28046*^ clones were obtained at 25°C by employing FLP/FRT mediated mitotic recombination with the following genotypes:

*eyFLP*/+; *mδ0.5-Gal4 FRT40/ubiGFP FRT40*; *UAS-mam*^*RNAi BL28046*^*aoslacZ*/+
*eyFLP/+; mδ0.5-Gal4 FRT40/ubiGFP FRT40; UAS-mam*^*RNAi BL28046*^/+

### Immunohistochemistry and Histology

Adult eye sectioning was performed as previously described (Jenny, 2011).

Third larval instar eye discs were dissected in ice-cold PBS and fixed in PBT (PBS+0.1% Triton-X)-4% formaldehyde for 12 minutes at room temperature. For immunohistochemistry, following primary antibodies were used: rat anti-DE-cad (1:20, DSHB), mouse anti-Fmi (1:10, DSHB), rabbit anti-β-gal (1:200, ICL), rat anti-Elav (1:100, DSHB), chicken anti-GFP (1:1000, Aves Labs). Secondary antibodies were obtained from Jackson Laboratories. Eye disc images were acquired by using Leica SP5 DMI microscope.

### Quantitative Analysis of Adult Eye Sections

The orientation of each ommatidium was marked based on the trapezoidal organization of the R-cells (see Figure 1B, D-H). A linear equator has been drawn along the boundary where two chiral forms meet. Clockwise and counter-clockwise angles from the equator to each ommatidia were measured for the black and red chiral forms respectively (see Figure 1B, D-H). Measurements were done by using ImageJ (National Institute of Health). The absolute values of measured angles from 3-4 independent eye sections for each genotype were pooled (300<n<550) and plotted in a polar histogram by using MATLAB. The angles were binned into 20° intervals between 0-180° and they were plotted in probability ratios from 0 to 1. For statistical analyses, the angles (α) were binned into 3 categories (α<60, 60<α<120, 120<α) for individual genotypes and chi-square test was performed.

### Quantitative Analysis of Eye Discs

The orientation of each ommatidium was marked perpendicular to the plane of R2/R5 cells (See Figure 1C). A linear equator was drawn perpendicular to the MF at the dorsoventral midline. Clockwise and counter-clockwise angles from the equator to each ommatidia were measured for the dorsal and ventral halves respectively. To avoid a potential bias due to the developmental delay in rotation from equator to the poles, the measurements were limited to the first 8 ommatidia from the equator for each row. Measurements were done by using ImageJ (National Institute of Health). The absolute values of measured angles from 7-8 independent eye discs (45<n<60) were pooled and violin plotted in PRISM. For statistical analyses, the angles (α) from individual rows were binned into 5 categories (α<40, 40<α<50, 50<α<60, 60<α70 and α>70) for each genotype and chi-square test was performed.

For *aos-lacZ* and Elav quantifications in *mδ0.5*>*mam*^*RNAi BL28046*^ mosaic eye discs, confocal stacks were maximum projected and individual cell intensities were measured in R3/R4 pairs between rows 5-11 by using ImageJ. Intensities in GFP^+^ and GFP^−^ R4 cells were normalized to their GFP^−^ R3 neighbors within each pair. Measurements from 5 discs were pooled (45<n<60) and plotted. For statistical analyses, the normalized intensity measurements (ι) were binned into 3 categories (ι<1, 1<ι<1.5, ι>1.5) for each genotype and chi-square test was performed.

For Fmi quantifications in *mδ0.5*>*mam*^*RNAi BL28046*^ mosaic eye discs, the confocal stacks were constructed in 3D and analyzed in IMARIS. Within GFP^+^ and GFP^−^ tissue, Fmi surface intensity on the apical membrane was measured for each R4 cell between rows 5-11 and normalized to the DE-cad intensity on the respective surface and plotted. Measurements from 5 discs were pooled (35<n<60) and plotted. For statistical analyses, the normalized intensity measurements (ι) were binned into 3 categories (ι<500, 500<ι<1000, ι>1000) for each genotype and *chi*-square test was performed.

## Acknowledgements

We thank the Bloomington Drosophila Research Center, Spyros Artavanis-Tsakonas and Utpal Banerjee for fly strains and reagents. We are grateful to all Mlodzik lab members for helpful input and discussions; and Robert Krauss, Cathie Pfleger, Timothy Blenkinsop and Jennifer Zallen for helpful comments and suggestions on the manuscript. Confocal laser scanning microscopy was performed at the ISMMS-Microscopy Core Facility supported by the Tisch Cancer Institute grant P30 CA196521 from the NCI. This work was supported by a NIH/NEI grant RO1 EY13256 to M.M.

## References

Ajuria, L., Nieva, C., Winkler, C., Kuo, D., Samper, N., Andreu, M.J., Helman, A., Gonzalez-Crespo, S., Paroush, Z., Courey, A.J., et al. (2011). Capicua DNA-binding sites are general response elements for RTK signaling in Drosophila. Development 138, 915–924.

Andersson, E.R., Sandberg, R., and Lendahl, U. (2011). Notch signaling: simplicity in design, versatility in function. Development 138, 3593–3612.

Baker, A.T., Zlobin, A., and Osipo, C. (2014). Notch-EGFR/HER2 Bidirectional Crosstalk in Breast Cancer. Front Oncol 4, 360.

Baker, N.E., Mlodzik, M., and Rubin, G.M. (1990). Spacing differentiation in the developing Drosophila eye: a fibrinogen-related lateral inhibitor encoded by scabrous. Science 250, 1370–1377.

Bao, S. (2014). Notch controls cell adhesion in the Drosophila eye. PLoS Genet 10, e1004087.

Baonza, A., and Freeman, M. (2001). Notch signalling and the initiation of neural development in the Drosophila eye. Development 128, 3889–3898.

Barolo, S., Stone, T., Bang, A.G., and Posakony, J.W. (2002). Default repression and Notch signaling: Hairless acts as an adaptor to recruit the corepressors Groucho and dCtBP to Suppressor of Hairless. Genes Dev 16, 1964–1976.

Blair, S.S. (1999). Eye development: Notch lends a handedness. Curr Biol 9, R356–360.

Brand, A.H., and Perrimon, N. (1993). Targeted gene expression as a means of altering cell fates and generating dominant phenotypes. Development 118, 401–415.

Bray, S.J. (2016). Notch signalling in context. Nat Rev Mol Cell Biol 17, 722–735.

Bray, S.J., and Gomez-Lamarca, M. (2018). Notch after cleavage. Curr Opin Cell Biol 51, 103–109.

Brou, C., Logeat, F., Gupta, N., Bessia, C., LeBail, O., Doedens, J.R., Cumano, A., Roux, P., Black, R.A., and Israel, A. (2000). A novel proteolytic cleavage involved in Notch signaling: the role of the disintegrin-metalloprotease TACE. Mol Cell 5, 207–216.

Brown, K.E., and Freeman, M. (2003). Egfr signalling defines a protective function for ommatidial orientation in the Drosophila eye. Development 130, 5401–5412.

Cagan, R.L., and Ready, D.F. (1989a). The emergence of order in the Drosophila pupal retina. Dev Biol 136, 346–362.

Cagan, R.L., and Ready, D.F. (1989b). Notch is required for successive cell decisions in the developing Drosophila retina. Genes Dev 3, 1099–1112.

Capilla, A., Johnson, R., Daniels, M., Benavente, M., Bray, S.J., and Galindo, M.I. (2012). Planar cell polarity controls directional Notch signaling in the Drosophila leg. Development 139, 2584–2593.

Choi, K.W., and Benzer, S. (1994). Rotation of photoreceptor clusters in the developing Drosophila eye requires the nemo gene. Cell 78, 125–136.

Chou, Y.H., and Chien, C.T. (2002). Scabrous controls ommatidial rotation in the Drosophila compound eye. Dev Cell 3, 839–850.

Cooper, M.T., and Bray, S.J. (1999). Frizzled regulation of Notch signalling polarizes cell fate in the Drosophila eye. Nature 397, 526–530.

Cooper, M.T., and Bray, S.J. (2000). R7 photoreceptor specification requires Notch activity. Curr Biol 10, 1507–1510.

Das, G., Reynolds-Kenneally, J., and Mlodzik, M. (2002). The atypical cadherin Flamingo links Frizzled and Notch signaling in planar polarity establishment in the Drosophila eye. Dev Cell 2, 655–666.

De Strooper, B., Annaert, W., Cupers, P., Saftig, P., Craessaerts, K., Mumm, J.S., Schroeter, E.H., Schrijvers, V., Wolfe, M.S., Ray, W.J., et al. (1999). A presenilin-1-dependent gamma-secretase-like protease mediates release of Notch intracellular domain. Nature 398, 518–522.

del Alamo, D., and Mlodzik, M. (2006). Frizzled/PCP-dependent asymmetric neuralized expression determines R3/R4 fates in the Drosophila eye. Dev Cell 11, 887–894.

Dobens, L.L., and Raftery, L.A. (2000). Integration of epithelial patterning and morphogenesis in Drosophila ovarian follicle cells. Dev Dyn 218, 80–93.

Dominguez, M., and de Celis, J.F. (1998). A dorsal/ventral boundary established by Notch controls growth and polarity in the Drosophila eye. Nature 396, 276–278.

Doroquez, D.B., and Rebay, I. (2006). Signal integration during development: mechanisms of EGFR and Notch pathway function and cross-talk. Crit Rev Biochem Mol Biol 41, 339–385.

Fanto, M., and Mlodzik, M. (1999). Asymmetric Notch activation specifies photoreceptors R3 and R4 and planar polarity in the Drosophila eye. Nature 397, 523–526.

Fiehler, R.W., and Wolff, T. (2007). Drosophila Myosin II, Zipper, is essential for ommatidial rotation. Dev Biol 310, 348–362.

Fiehler, R.W., and Wolff, T. (2008). Nemo is required in a subset of photoreceptors to regulate the speed of ommatidial rotation. Dev Biol 313, 533–544.

Flores, G.V., Duan, H., Yan, H., Nagaraj, R., Fu, W., Zou, Y., Noll, M., and Banerjee, U. (2000). Combinatorial signaling in the specification of unique cell fates. Cell 103, 75–85.

Furriols, M., and Bray, S. (2000). Dissecting the mechanisms of suppressor of hairless function. Dev Biol 227, 520–532.

Gaengel, K., and Mlodzik, M. (2003). Egfr signaling regulates ommatidial rotation and cell motility in the Drosophila eye via MAPK/Pnt signaling and the Ras effector Canoe/AF6. Development 130, 5413–5423.

Guruharsha, K.G., Kankel, M.W., and Artavanis-Tsakonas, S. (2012). The Notch signalling system: recent insights into the complexity of a conserved pathway. Nat Rev Genet 13, 654–666.

Haruki, N., Kawaguchi, K.S., Eichenberger, S., Massion, P.P., Olson, S., Gonzalez, A., Carbone, D.P., and Dang, T.P. (2005). Dominant-negative Notch3 receptor inhibits mitogen-activated protein kinase pathway and the growth of human lung cancers. Cancer Res 65, 3555–3561.

Hasson, P., and Paroush, Z. (2006). Crosstalk between the EGFR and other signalling pathways at the level of the global transcriptional corepressor Groucho/TLE. Br J Cancer 94, 771–775.

Henrique, D., and Schweisguth, F. (2019). Mechanisms of Notch signaling: a simple logic deployed in time and space. Development 146.

Housden, B.E., Terriente-Felix, A., and Bray, S.J. (2014). Context-dependent enhancer selection confers alternate modes of notch regulation on argos. Mol Cell Biol 34, 664–672.

Jenny, A. (2010). Planar cell polarity signaling in the Drosophila eye. Curr Top Dev Biol 93, 189–227.

Jenny, A., Darken, R.S., Wilson, P.A., and Mlodzik, M. (2003). Prickle and Strabismus form a functional complex to generate a correct axis during planar cell polarity signaling. EMBO J 22, 4409–4420.

Kenyon, K.L., Ranade, S.S., Curtiss, J., Mlodzik, M., and Pignoni, F. (2003). Coordinating proliferation and tissue specification to promote regional identity in the Drosophila head. Dev Cell 5, 403–414.

Klein, D.E., Nappi, V.M., Reeves, G.T., Shvartsman, S.Y., and Lemmon, M.A. (2004). Argos inhibits epidermal growth factor receptor signalling by ligand sequestration. Nature 430, 1040–1044.

Klein, D.E., Stayrook, S.E., Shi, F., Narayan, K., and Lemmon, M.A. (2008). Structural basis for EGFR ligand sequestration by Argos. Nature 453, 1271–1275.

Kopan, R., and Ilagan, M.X. (2009). The canonical Notch signaling pathway: unfolding the activation mechanism. Cell 137, 216–233.

Kozlovskaja-Gumbriene, A., Yi, R., Alexander, R., Aman, A., Jiskra, R., Nagelberg, D., Knaut, H., McClain, M., and Piotrowski, T. (2017). Proliferation-independent regulation of organ size by Fgf/Notch signaling. Elife 6.

Krejci, A., Bernard, F., Housden, B.E., Collins, S., and Bray, S.J. (2009). Direct response to Notch activation: signaling crosstalk and incoherent logic. Sci Signal 2, ra1.

Lieber, T., Kidd, S., and Young, M.W. (2002). kuzbanian-mediated cleavage of Drosophila Notch. Genes Dev 16, 209–221.

Markopoulou, K., Welshons, W.J., and Artavanis-Tsakonas, S. (1989). Phenotypic and molecular analysis of the facets, a group of intronic mutations at the Notch locus of Drosophila melanogaster which affect postembryonic development. Genetics 122, 417–428.

McNeill, H., Craig, G.M., and Bateman, J.M. (2008). Regulation of neurogenesis and epidermal growth factor receptor signaling by the insulin receptor/target of rapamycin pathway in Drosophila. Genetics 179, 843–853.

Mirkovic, I., Gault, W.J., Rahnama, M., Jenny, A., Gaengel, K., Bessette, D., Gottardi, C.J., Verheyen, E.M., and Mlodzik, M. (2011). Nemo kinase phosphorylates beta-catenin to promote ommatidial rotation and connects core PCP factors to E-cadherin-beta-catenin. Nat Struct Mol Biol 18, 665–672.

Mirkovic, I., and Mlodzik, M. (2006). Cooperative activities of drosophila DE-cadherin and DN-cadherin regulate the cell motility process of ommatidial rotation. Development 133, 3283–3293.

Mlodzik, M. (1999). Planar polarity in the Drosophila eye: a multifaceted view of signaling specificity and cross-talk. EMBO J 18, 6873–6879.

Morel, V., Lecourtois, M., Massiani, O., Maier, D., Preiss, A., and Schweisguth, F. (2001). Transcriptional repression by suppressor of hairless involves the binding of a hairless-dCtBP complex in Drosophila. Curr Biol 11, 789–792.

Mumm, J.S., Schroeter, E.H., Saxena, M.T., Griesemer, A., Tian, X., Pan, D.J., Ray, W.J., and Kopan, R. (2000). A ligand-induced extracellular cleavage regulates gamma-secretase-like proteolytic activation of Notch1. Mol Cell 5, 197–206.

Papayannopoulos, V., Tomlinson, A., Panin, V.M., Rauskolb, C., and Irvine, K.D. (1998). Dorsal-ventral signaling in the Drosophila eye. Science 281, 2031–2034.

Pezeron, G., Millen, K., Boukhatmi, H., and Bray, S. (2014). Notch directly regulates the cell morphogenesis genes Reck, talin and trio in adult muscle progenitors. J Cell Sci 127, 4634–4644.

Polacheck, W.J., Kutys, M.L., Yang, J., Eyckmans, J., Wu, Y., Vasavada, H., Hirschi, K.K., and Chen, C.S. (2017). A non-canonical Notch complex regulates adherens junctions and vascular barrier function. Nature 552, 258–262.

Rebay, I., Fleming, R.J., Fehon, R.G., Cherbas, L., Cherbas, P., and Artavanis-Tsakonas, S. (1991). Specific EGF repeats of Notch mediate interactions with Delta and Serrate: implications for Notch as a multifunctional receptor. Cell 67, 687–699.

Reynolds-Kenneally, J., and Mlodzik, M. (2005). Notch signaling controls proliferation through cell-autonomous and non-autonomous mechanisms in the Drosophila eye. Dev Biol 285, 38–48.

Rohrbaugh, M., Ramos, E., Nguyen, D., Price, M., Wen, Y., and Lai, Z.C. (2002). Notch activation of yan expression is antagonized by RTK/pointed signaling in the Drosophila eye. Curr Biol 12, 576–581.

Roignant, J.Y., and Treisman, J.E. (2009). Pattern formation in the Drosophila eye disc. Int J Dev Biol 53, 795–804.

Struhl, G., and Adachi, A. (1998). Nuclear access and action of notch in vivo. Cell 93, 649–660.

Struhl, G., and Greenwald, I. (1999). Presenilin is required for activity and nuclear access of Notch in Drosophila. Nature 398, 522–525.

Strutt, H., and Strutt, D. (1999). Polarity determination in the Drosophila eye. Curr Opin Genet Dev 9, 442–446.

Strutt, H., and Strutt, D. (2003). EGF signaling and ommatidial rotation in the Drosophila eye. Curr Biol 13, 1451–1457.

Tomlinson, A., and Ready, D.F. (1987). Neuronal differentiation in Drosophila ommatidium. Dev Biol 120, 366–376.

Tomlinson, A., and Struhl, G. (1999). Decoding vectorial information from a gradient: sequential roles of the receptors Frizzled and Notch in establishing planar polarity in the Drosophila eye. Development 126, 5725–5738.

Tomlinson, A., and Struhl, G. (2001). Delta/Notch and Boss/Sevenless signals act combinatorially to specify the Drosophila R7 photoreceptor. Mol Cell 7, 487–495.

Torres, I.L., Lopez-Schier, H., and St Johnston, D. (2003). A Notch/Delta-dependent relay mechanism establishes anterior-posterior polarity in Drosophila. Dev Cell 5, 547–558.

Vinos, J., and Freeman, M. (2000). Evidence that Argos is an antagonistic ligand of the EGF receptor. Oncogene 19, 3560–3562.

Weber, U., Paricio, N., and Mlodzik, M. (2000). Jun mediates Frizzled-induced R3/R4 cell fate distinction and planar polarity determination in the Drosophila eye. Development 127, 3619–3629.

Weber, U., Pataki, C., Mihaly, J., and Mlodzik, M. (2008). Combinatorial signaling by the Frizzled/PCP and Egfr pathways during planar cell polarity establishment in the Drosophila eye. Dev Biol 316, 110–123.

Wilson, J.J., and Kovall, R.A. (2006). Crystal structure of the CSL-Notch-Mastermind ternary complex bound to DNA. Cell 124, 985–996.

Winter, C.G., Wang, B., Ballew, A., Royou, A., Karess, R., Axelrod, J.D., and Luo, L. (2001). Drosophila Rho-associated kinase (Drok) links Frizzled-mediated planar cell polarity signaling to the actin cytoskeleton. Cell 105, 81–91.

Wolff, T., and Ready, D.F. (1991). The beginning of pattern formation in the Drosophila compound eye: the morphogenetic furrow and the second mitotic wave. Development 113, 841–850.

Wolff, T., and Rubin, G.M. (1998). Strabismus, a novel gene that regulates tissue polarity and cell fate decisions in Drosophila. Development 125, 1149–1159.

Wu, J., and Mlodzik, M. (2009). A quest for the mechanism regulating global planar cell polarity of tissues. Trends Cell Biol 19, 295–305.

Wu, L., Aster, J.C., Blacklow, S.C., Lake, R., Artavanis-Tsakonas, S., and Griffin, J.D. (2000). MAML1, a human homologue of Drosophila mastermind, is a transcriptional co-activator for NOTCH receptors. Nat Genet 26, 484–489.

Yan, H., Chin, M.L., Horvath, E.A., Kane, E.A., and Pfleger, C.M. (2009). Impairment of ubiquitylation by mutation in Drosophila E1 promotes both cell-autonomous and non-cell-autonomous Ras-ERK activation in vivo. J Cell Sci 122, 1461–1470.

Yoo, A.S., Bais, C., and Greenwald, I. (2004). Crosstalk between the EGFR and LIN-12/Notch pathways in C. elegans vulval development. Science 303, 663–666.

Zacharioudaki, E., and Bray, S.J. (2014). Tools and methods for studying Notch signaling in Drosophila melanogaster. Methods 68, 173–182.

Zacharioudaki, E., Housden, B.E., Garinis, G., Stojnic, R., Delidakis, C., and Bray, S.J. (2016). Genes implicated in stem cell identity and temporal programme are directly targeted by Notch in neuroblast tumours. Development 143, 219–231.

Zheng, L., Zhang, J., and Carthew, R.W. (1995). frizzled regulates mirror-symmetric pattern formation in the Drosophila eye. Development 121, 3045–3055.

