## Supplemental Data for "Notch signaling coordinates ommatidial rotation in the Drosophila eye via transcriptional regulation of the EGF-Receptor ligand Argos"

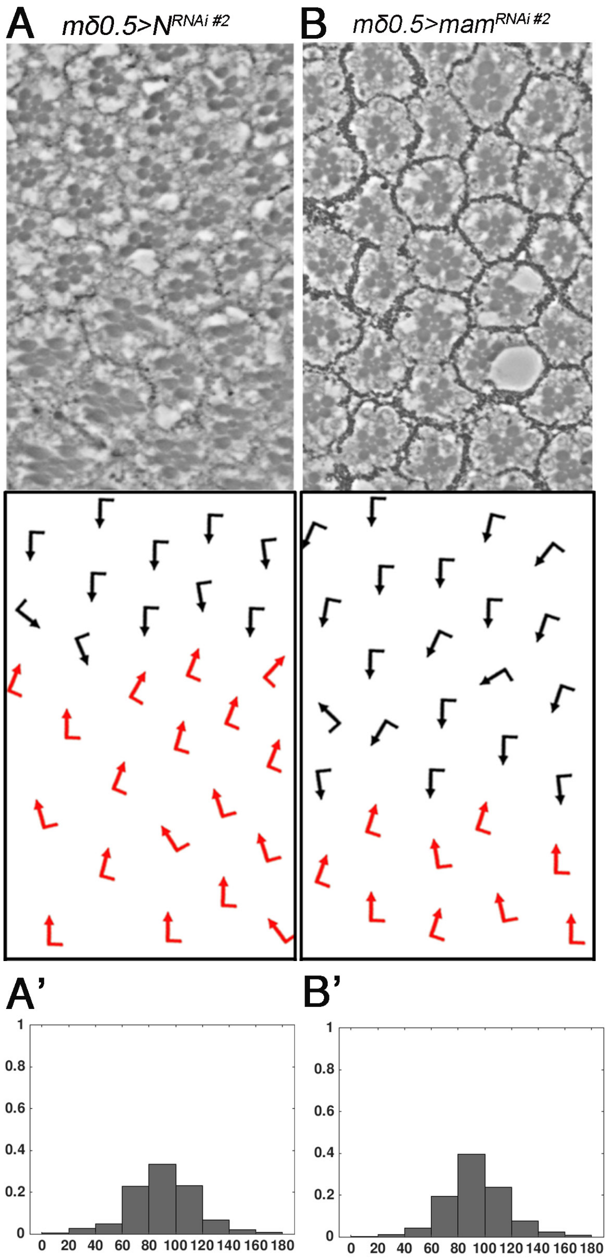


**Figure S1. Perturbation of Notch signaling in the eye leads to misorientation of ommatidia.** Adult eye sections with ommatidial orientation schematics and ommatidial orientation angle histograms for *mδ0.5>N*$^{RNAi\ \#2}$ (*BL7078*) **(A, A')** and *mδ0.5>mam*$^{RNAi\ \#2}$ (*BL63601)* **(B, B'),** n>300, 3 eyes each.

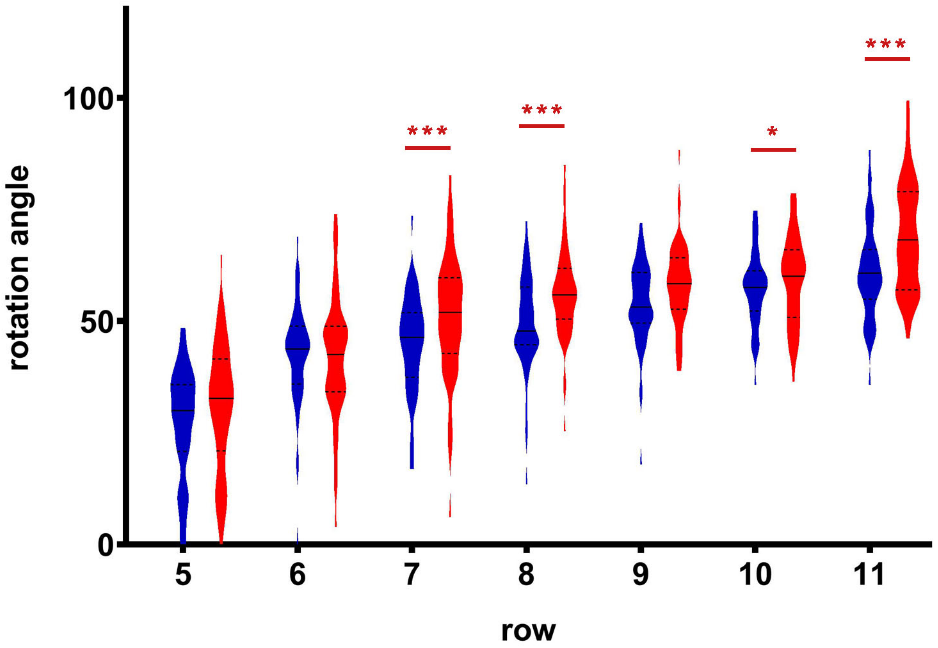


**Figure S2. Notch signaling is required in R3/R4 pairs to regulate OR.** Quantification of rotation angles observed in individual preclusters in rows 5-11 as plotted for control (wild type) in blue*;* *mδ0.5>mam*$^{RNAi}$ (*BL63601*) in green. Statistical analyses were performed for each row between genotypes. Asterisks denote significance by chi-square test (* p<0.05, ** p<0.005, *** p<0.0005).

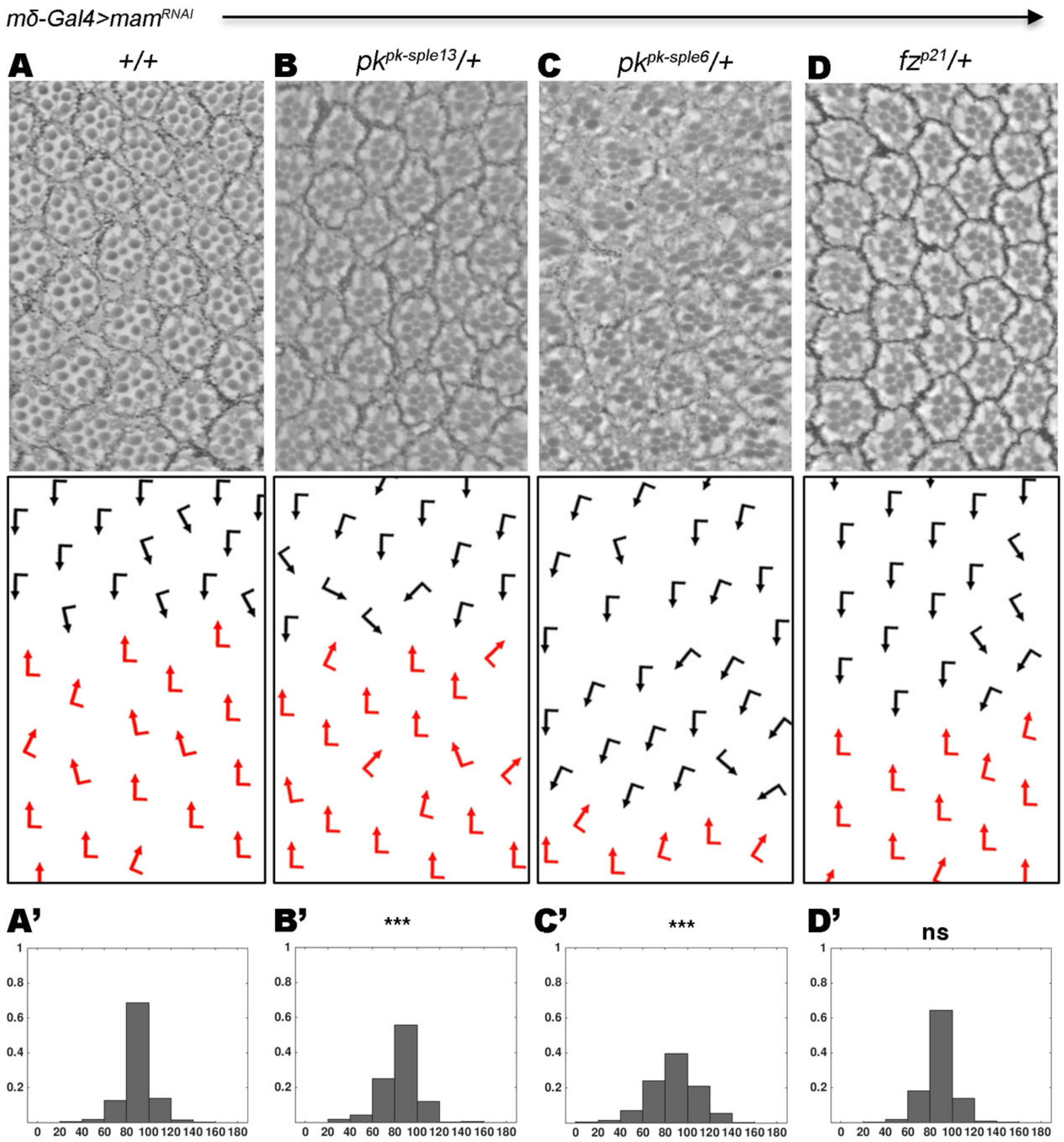


**Figure S3. *mam* genetically interacts with *pk* to regulate OR.**

**(A-D')** Adult eye sections with ommatidial orientation schematics and ommatidial orientation angle histograms of eyes from *mδ0.5Gal4>mam$^{RNAi}$ (BL28046)* in the following genetic backgrounds: **(A-A')** +/+ (*wt* control); **(B-B')** *pk$^{pk\text{-}sple13}$*/+, **(C-C')** *pk$^{pk\text{-}sple6}$*/+ and **(D-D')** *fz$^{p21}$*/+. Asterisks denote significance by chi-square test (*** p<0.0005). Note robust enhancement of the *mδ0.5Gal4>mam$^{RNAi}$* rotation phenotype by both *pk$^{-/+}$* genotypes.

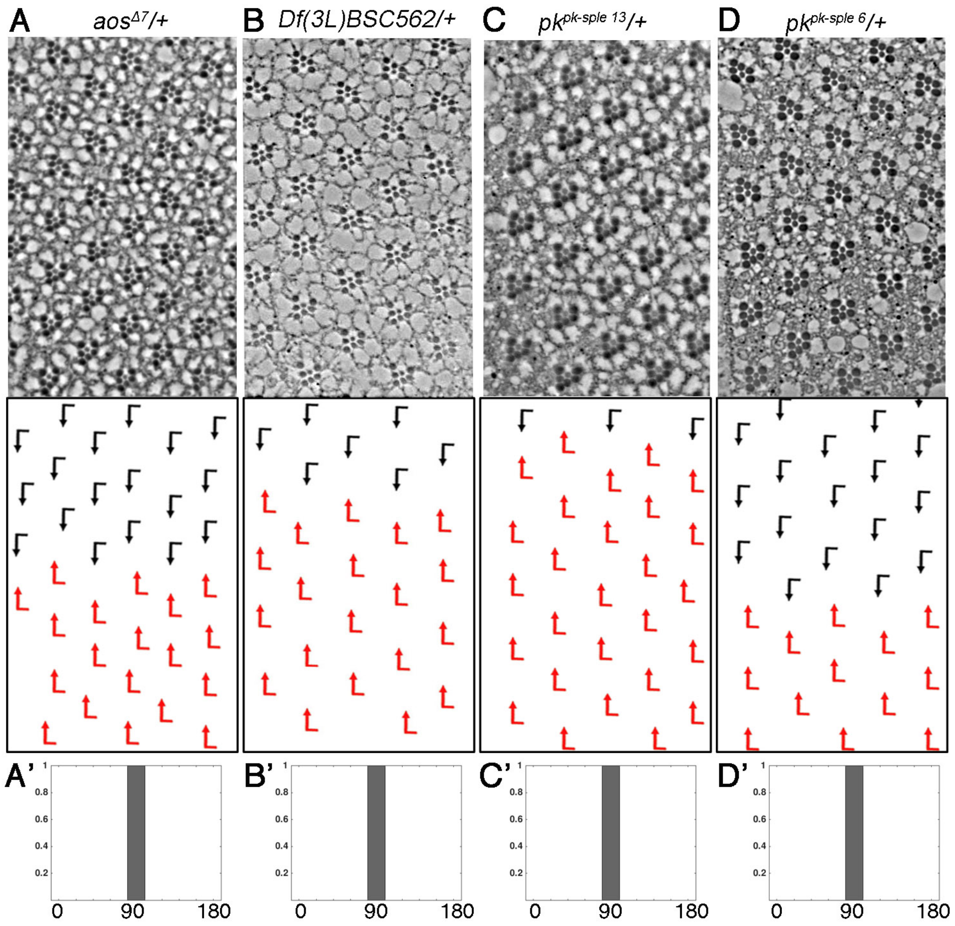


**Figure S4. Heterozygous mutations in *aos* and *pk* that interact with the Notch-signaling genotypes do not show dominant effects in an otherwise wild-type background.**

**(A-D')** Adult eye sections with ommatidial orientation schematics and ommatidial orientation angle histograms of eyes from the following genetic backgrounds: **(A-A')** *aosΔ7/+* ; **(B-B')** *DF(3L)BSC562/+*, **(C-C')** $pk^{pk\text{-}sple13}$/+, and **(D-D')** $pk^{pk\text{-}sple6}$/+. Note that all eyes have wild-type appearance in these heterozygous backgrounds.

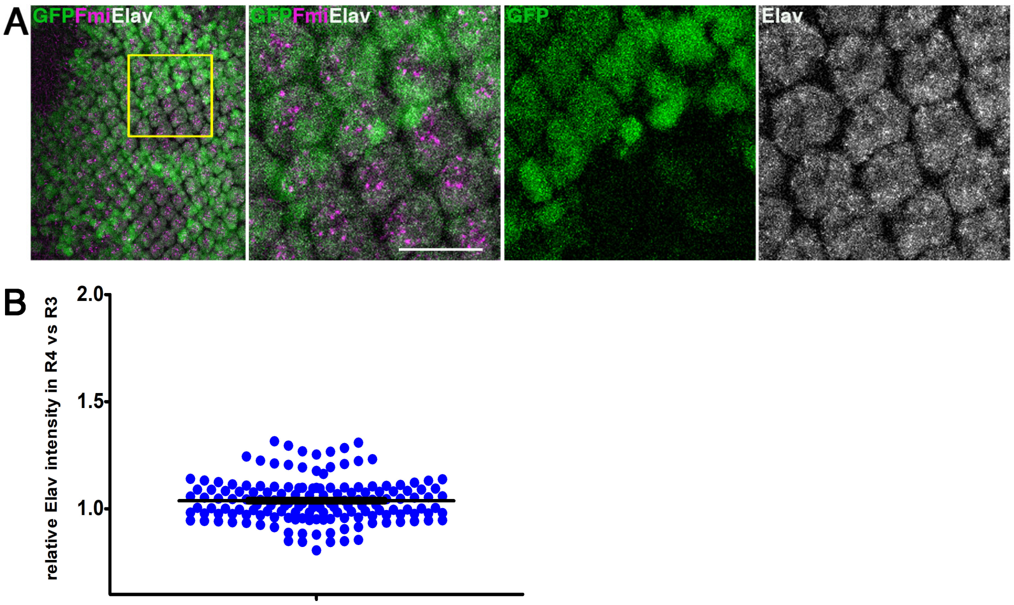


**Figure S5. Notch signaling does not affect Elav levels in R4. (A)** Third larval instar eye imaginal discs mosaic for *mδ0.5>mam*$^{RNAi}$ *(BL28046*; marked by absence of GFP/green) stained for Fmi (magenta) and Elav (gray). Note that the level of Elav is indistinguishable in the wild-type (GFP, green) vs the mutant clonal area (black). See Figure 4F in main text for comparison to *aos* expression. Scale bar is 10 μm. **(B)** Quantification of relative intensity of Elav in R4 vs. R3, note that the ratio is basically 1, meaning that there is no differential expression between the two cells.

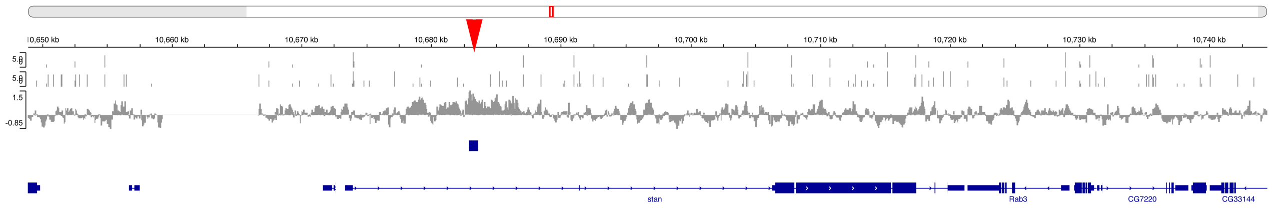


**Figure S6. Occupation by Su(H) in genomic sequences of the *fmi/stan* locus.**

ChIP enrichment for Su(H)-occupancy at the *stan/fmi* locus in CNS samples ($\alpha$-Su(H) enrichment relative to input, scale $\log_2$) (Zacharioudaki et al., 2016). Blue bar indicates potential the region of significant enrichment. Gray bars indicate the positions of Su(H)-binding motifs; bar height represents the motif-score (scale 0-5); upper graph indicates motif conservation across 12 *Drosophila* species. Gene regions are depicted in dark blue. Although some enrichment is detected (small blue bar above large intron in gene schematic), there are no conserved binding site in that region (red arrowhead), suggesting that it is not meaningful. These data are consistent with the notion that Notch-Su(H) signaling is not a transcriptional regulator of *fmi/stan.* Furthermore, there was no ChIP enrichment detected associated with genomic regions of the *pk* locus.
